# Novel Roles of Sonic Hedgehog Signaling in Retinal Patterning and Neurogenesis During Mammalian Eye Development

**DOI:** 10.1101/2025.08.19.671114

**Authors:** Miranda R. Krueger, Simranjeet K. Cheema, Sergi Simo, Edward M. Levine, Nadean L. Brown, Anna La Torre

**Affiliations:** Department of Cell Biology and Human Anatomy. School of Medicine. University of California, Davis, United States; Vanderbilt Eye Institute, Vanderbilt University Medical Center, Nashville, United States

## Abstract

The Sonic Hedgehog (Shh) signaling pathway is essential for the patterning, growth, and morphogenesis of many tissues. During early eye development, Shh is critical for the formation of the two optic vesicles, which give rise to the retina, retinal pigment epithelium (RPE), and optic stalk. It also regulates the balance between cell proliferation and differentiation during retinal histogenesis, a key process in shaping the cellular architecture of the mature retina. Despite these well-established roles, the temporal dynamics, region-specific functions, and downstream consequences of Shh signaling during retinal development remain poorly understood. Here, we present a comprehensive analysis of Shh signaling across multiple stages of retinal development using temporally and spatially controlled deletion of Smoothened (Smo), an essential transducer of the pathway. This approach reveals previously unrecognized requirements for Shh signaling in specifying optic nerve head identity and maintaining nasal-temporal polarity. We also show that Shh signaling coordinates neurogenesis by sustaining the retinal progenitor pool while also regulating progenitor competence, ensuring appropriate proportions of retinal cell types. Our data indicate that both proliferative capacity and the timing of cell fate specification are shaped by Shh pathway activity. Together, these findings establish new mechanistic links between Shh signaling, regional patterning, and temporal regulation of neurogenesis, providing novel insights into how morphogen signaling is repurposed across developmental time to orchestrate complex tissue architecture.

## INTRODUCTION

The vertebrate eye is a complex organ primarily composed of the retina, optic nerve, retinal pigment epithelium (RPE), iris, ciliary body, and lens. Each of these components fulfills distinct but essential functions. Accordingly, all these tissues need to be assembled in an exquisitely precise three-dimensional organization that is achieved during development (Fuhrmann, 2010, Casey et al., 2023, Zhang et al., 2023b).

The development of the eye begins when a patch of anterior neural plate cells, known as the eye field, is specified by a conserved gene regulatory network (Zuber et al., 2003), which in mice occurs around embryonic day 7 (E7). Influenced by different signaling molecules, including Sonic Hedgehog (Shh) (Chiang et al., 1996, Ohtsuka et al., 2022), the eye field separates into bilateral optic vesicles that evaginate from the ventral diencephalon around E8. These vesicles grow laterally and are subsequently triggered by the surface ectoderm to invaginate into optic cups, but remain connected to the brain through the optic stalks. Each optic cup consists of an outer layer that will become the non-neuronal RPE and an inner layer that will differentiate into the neural retina. The optic cup is patterned along its dorsal-ventral, nasal-temporal, and proximo-distal axes by multiple extrinsic factors (Stull and Wikler, 2000, Sakuta et al., 2001, Murali et al., 2005, Picker and Brand, 2005, Kobayashi et al., 2010). These patterning processes produce specialized retinal subdomains such as the optic nerve head (ONH) and the ciliary marginal zone (CMZ) (Torres et al., 1996, Reh and Levine, 1998, Marcucci et al., 2016).

Between E11-12 mouse retinal progenitor cells (RPCs) begin to exit cell cycle and differentiate following a conserved sequence. Retinal neurogenesis begins with the production of retinal ganglion cells (RGCs), cone photoreceptors, horizontal cells, and GABAergic amacrine cells, including the starburst subtype, that appear at early stages of development, while glycinergic amacrine cells, rod photoreceptors, bipolar cells, and Müller glia are generated later (Livesey and Cepko, 2001, Rapaport et al., 2004, Wallace, 2011, Clark et al., 2019).

All these intricate developmental processes are essential for establishing the proper architecture and functionality of the eye, ultimately enabling vision. Disruptions during early specification and patterning can lead to congenital eye malformations such as anophthalmia, microphthalmia, coloboma, or optic nerve hypoplasia (Cavodeassi et al., 2019).

The formation and patterning of the different regions and cells within the optic cup involve a complex network of signaling pathways and molecules, such as Shh, Notch, Bone Morphogenetic proteins (BMPs), Fibroblast Growth Factors (FGFs), and Retinoic Acid (RA) (Levine et al., 1997, Hyer et al., 1998, Vogel-Hopker et al., 2000, Zhao et al., 2001, Zhang and Yang, 2001b, Perron et al., 2003, Esteve and Bovolenta, 2006, Morcillo et al., 2006, Duester, 2009, Riesenberg et al., 2009, Zhao et al., 2010, Bosze et al., 2023). However, the precise roles of these pathways at different stages of development, as well as their interactions, are not yet fully understood.

Shh is a secreted glycoprotein initially expressed by the ventral midline of the neural tube (Ericson et al., 1995, Dale et al., 1997). Upon binding to the Patched1 (Ptch1) receptor on target cells, inhibition of the GPCR transmembrane protein Smoothened (Smo) is relieved (Taipale et al., 2002, Lum and Beachy, 2004). Downstream of Smo, a multimolecular network transduces Shh signaling that ultimately modifies the activity of Gli proteins. Accordingly, in the absence of Shh, Ptch1 inhibits Smo, and Gli proteins undergo proteolytic processing, resulting in truncated, repressive forms (Hui and Angers, 2011). Conversely, when Shh signaling is active, full-length Gli proteins accumulate in the nucleus, promoting the transcription of key target genes, including *Gli1* and *Ptch1* (Bonifas et al., 2001).

Widespread loss of Shh signaling at early stages of development results in severe morphological and structural abnormalities, including the failure of the anterior neural plate to properly split into symmetrical bilateral regions, resulting in cyclopia (Chiang et al., 1996) and holoprosencephaly (Roessler et al., 1996). Previous studies have also implicated Shh in the establishment of spatial axes. In chicken and mouse embryos, BMP and Shh signaling exhibit antagonistic effects through the dorsal determinant *Tbx5* and its ventral counterpart *Vax2* to establish dorsal and ventral fates of the optic cup, respectively (Zhang and Yang, 2001b). However, the full extent of Shh function in organizing the regional patterning of the developing eye remains incompletely understood.

Here, we investigate the distinct roles of Shh signaling across multiple stages of eye development by generating Shh loss-of-function models using *Smo* conditional mutant mice and leveraging the activities of different Cre drivers to precisely control the timing and location of loss of *Smo* activity. Our data reveal an underappreciated role this pathway in establishing optic nerve head (ONH) cell identity, particularly in the naso-ventral quadrant of the optic cup. Similarly, we demonstrate a previously unrecognized function for Shh in naso-temporal retinal patterning.

Previous studies have shown that Shh is expressed by RGCs, acts on RPCs, and is essential for the proper control of RPC proliferation (Wang et al., 2002, Dakubo et al., 2003, Wall et al., 2009). However, the extent to which Shh roles influences the specification of the other major retinal cell classes has not been reported (Zhang and Yang, 2001a, Wang et al., 2005, Sakagami et al., 2009). Here, we show that *Smo* ablation disrupts the normal balance of all retinal cell populations through both precocious cell cycle exit and misregulation of progenitor competence timing.

Together, these findings underscore the critical and multifaceted role of Shh in shaping retinal architecture. Our study advances our understanding of the molecular mechanisms guiding spatial and temporal organization in the developing eye, and provides a foundation for investigating how disruptions in Shh signaling contribute to congenital ocular disorders.

## MATERIALS AND METHODS

### Animals

Animals were handled with approval from the University of California Davis Institutional Animal Care and Use Committees (IACUC) and were housed and cared for in accordance with the guidelines provided by the National Institutes of Health.

*Smo* conditional mice were obtained from JAX (Smo^tm2Amc^; JAX stock number 004526). Rax-Cre BAC transgenic line Tg(Rax-cre) NL44Gsat/Mmucd was created by the GENSAT project and cryobanked at MMRRC UC Davis (stock number 034748-UCD, (Bosze et al., 2020)). Chx10-Cre was generated by (Rowan and Cepko, 2004) *Tg(Chx10-EGFP/cre;-ALPP)2Clc*; JAX stock number 005105. All lines were maintained on a CD1 background. PCR genotyping was performed per JAX or MMRRC protocols. Conditional mutant breeding schemes mated one Cre Tg/+ mouse to another homozygous for the Smo conditional allele to create Cre; Smo^CKO/+^ mice, which were used in timed matings to Smo^CKO/CKO^ mice. The date a vaginal plug was noted, was assigned the age E0.5.

### Sample collection and preparation

Embryos and postnatal eyes were collected in cold PBS, fixed and prepared for cryo-embedding or paraffin-embedding. For cryopreservation, samples were submerged in 4% paraformaldehyde/PBS for 1 hour or overnight at 4°C, cryoprotected through a stepwise sucrose gradient, embedded and frozen in Optimum Cutting Temperature (OCT) compound (Tissue-Tek). For paraffin embedding, samples were fixed in modified Carnoy’s fixative (60% ethanol, 30% formaldehye, and 10% acetic acid) overnight at 4°C, dehydrated through an ethanol/water gradient and xylene, then embedded into paraffin blocks. Retinas were sectioned (15μm for cryo or 6μm paraffin) on a horizontal (transverse), coronal, or sagittal plane as specified in each figure.

### Immunohistochemistry and RNAscope in situ hybridization

Cryosections were rinsed with PBS to remove all OCT and permeabilized with 0.3% Triton X-100/PBS. Paraffin sections were deparaffinized using xylene, rehydrated through an ethanol gradient, permeabilized in 0.3% Triton X-100/PBS, then antigen unmasked in 95°C 0.01 M sodium citrate pH 8 twice for 5 min. Sections were blocked with 10% normal donkey serum/0.1% Triton X-100/PBS for 1 hr at room temperature. Primary antibodies were diluted in blocking solution for incubation overnight at 4°C. See Table 1 for antibody details and concentrations used. Species-specific, fluorescently labeled secondary antibodies were diluted in blocking solution with 4′,6-diamidino-2-phenylindole (DAPI) (Sigma-Aldrich) for nuclear counter-staining for 1hr at RT. Slides were mounted with Fluoromount-G (Southern Biotech). For RNAscope detection, protocols were performed according to the RNAscope Multiplex Fluorescent Reagent Kit v2 Assay manual using the provided reagents and as described before (Krueger et al., 2024). Briefly, following sample fixation and preparation described above, paraffin slides were baked at 60°C and deparaffined. Sections were then pretreated with 5% hydrogen peroxide at RT for 10 min, target antigen retrieval for 20 min at 99°C, and protease plus for 30 min at 40°C. For cryosections, samples were rinsed with 0.3% Triton X-100/PBS, quick pretreatment with hydrogen peroxide, target retrieval for 5 min at 99°C, rinsed with 100% EtOH, and baked for 5 min at 60°C. All sections were then incubated with the appropriate hybridization probes for 2 hr at 40°C. Probes include: Smo (Ref:577901-C3), Ptch1 (Ref:402811-C2), Gli1 (Ref:311001-C3), Foxd1 (Ref:495501-C3), Cdk6 (Ref:570091-C1), and Dio3 (Ref:561641-C2). Images were obtained by Fluoview FV4000 confocal microscope (Olympus) or Axio Imager.M2 with Apotome.2 microscope system (Zeiss). All figures were assembled using Photoshop and Illustrator (Adobe).

**Table 1:** Primary Antibodies.

| **Antibody** | **Source** | **Catalog** | **Concentration** |
| --- | --- | --- | --- |
| Anti-Arl13b (Rabbit) | Proteintech | 17711-1AP | 1:1000 |
| Anti-Atoh7 (Rabbit) | Novus | NBP1-88639 | 1:500 |
| Anti-AP2-alpha (Mouse) | Thermofisher | 3B5 | 1:200 |
| Anti-Boc (Goat) | R&D Systems | AF2385 | 1:500 |
| Anti-BRN3 (Goat) | Santa Cruz | sc-6026 | 1:100 |
| Anti-Calbindin (Mouse) | Neuromab | 73-452 | 1:250 |
| Anti-Calretinin (Goat) | Swat | CG1 | 1:200 |
| Anti-Cdo (Goat) | R&D Systems | AF2429 | 1:500 |
| Anti-Cone Arrestin<br>(Rabbit) | Millipore | Ab15282 | 1:400 |
| Anti-CRX<br>(Mouse) | Abnova | H00001406-M02 | 1:200 |
| Anti-FoxG1 (Rabbit) | Abcam | Ab196868 | 1:200 |
| Anti-Gas1 (Goat) | R&D Systems | AF2644 | 1:500 |
| Anti-GFAP<br>(Mouse) | Sigma | 0000124890 | 1:100 |
| Anti-Glutamine Synthetase (Mouse) | Millipore | MAB302 | 1:100 |
| Ib4 lectin | Life Technologies | I21411 | 1:100 |
| Anti-Laminin (Rabbit) | Abcam | AB11575 | 1:500 |
| Anti-Lrp2 (Rabbit) | Abcam | AB76969 | 1:1000 |
| Anti-Op sin Red/Green (Rabbit) | Millipore | AB5405 | 1:200 |
| Anti-Op sin Blue<br>(Goat) | Santa Cruz | sc-14363 | 1:200 |
| Anti-OTX2<br>(Goat) | R&D Systems | AF1979 | 1:200 |
| Anti-PAX2<br>(Rabbit) | Biolegend | 901001 | 1:1000 |
| Anti-PAX6<br>(Mouse) | Santa Cruz | sc-32766 | 1:50 |
| Anti-PCNA (Mouse) | Invitrogen | 13-3900 | 1:200 |
| Anti-PH3<br>(Rabbit) | Thermo Fisher | PA5-17869 | 1:100 |
| Anti-RBPMS (Rabbit) | Proteintech | 15187-1-AP | 1:400 |
| Anti-SOX2<br>(Goat) | R&D Systems | AF2018 | 1:100 |
| Anti-SOX9<br>(Rabbit) | Millipore | AB5535 | 1:200 |
| Anti-TUBB3<br>(Mouse) | BioLegend | 801201 | 1:500 |

### Hematoxylin and eosin (H&E) staining

Sections were deparaffinized using xylene, rehydrated with stepwise ethanol/water solutions, stained with hematoxylin for 7 min, quickly rinsed with acid alcohol and ammonia water for 15 secs, counterstained with eosin for 6 min, and dehydrated in ethanol/water gradient of solutions. Sections were washed using xylene and mounted for microscopy using xylene diluted Permount (Fisher Chemical).

### Statistical analyses

The number of biological replicates and the statistical analyses employed are detailed in the corresponding figures and figure legends. Data presented as mean ± SEM. Retinal length and diameter were quantified for each side of the retina (temporal and nasal) starting from the ONH to the ciliary margin across three biological replicates. Similarly, retinal layer thickness quantifications were obtained at equidistant areas from the ONH across all samples. For cell-type histological quantifications, single or double fluorescently labelled cells were calculated across at least 3 biological replicates and normalized per area. For all pairwise analysis to obtain mean and p-values, student t-tests were performed. ANOVA analyses were used to compare across multiple groups. Statistical significance, p-values<0.05 were considered significant. All statistical analyses and plot generation was performed using Prism 9 (GraphPad).

### Flat-mount preparations

Eyes were enucleated from postnatal day 21 (P21) mice and fixed in 4% PFA/PBS for 30 minutes at room temperature prior to retina dissection. Isolated retinas were post-fixed in 4% PFA overnight at 4°C, followed by tissue permeabilization in 1% Triton X-100 in PBS for 1 hour at room temperature. Samples were blocked in 10% normal donkey serum/1% Triton X-100 in PBS supplemented for 2 hours at room temperature, the incubated with primary antibodies diluted in this blocking solution for 72 hours at 4°C, washed three times in PBS, then incubated in diluted secondary antibodies overnight at 4°C, and flat-mounted using Fluoromount-G (SouthernBiotech).

### RNAseq

Total RNA from control (*n*=4) and Rax-Cre;SmoCKO (*n*=5) isolated retinas was extracted using the Total RNA Purification Plus Kit (Norgen Biotek Corp., 48300). Bulk RNA sequencing was carried by the DNA Technologies and Expression Analysis Core at the UC Davis Genome Center, supported by NIH Shared Instrumentation Grant 1S10OD010786-01. Libraries were generated for 3’-Tag-Seq Gene Expression Profiling at 4M reads per sample. The fragment size distribution was verified using micro-capillary gel electrophoresis on a Bioanalyzer 2100 (Agilent). These libraries were then sent for quality control using fluorometry on a Qubit instrument (Life Technologies). Finally, 3’-Tag-Seq libraries were sequenced via single-end sequencing on the HiSeq 4000 (Illumina). Raw sequencing reads were assessed for quality. Adapters and low-quality bases were trimmed and cleaned reads were then aligned to the mouse reference genome. Gene-level counts were quantified using featureCounts from the Subread package, based on the corresponding gene annotation file (Ensembl). Differential gene expression analysis was performed. Genes with an adjusted p-value < 0.05 and absolute log2 fold change > 1 were considered significantly differentially expressed.

### EdU Birthdating

Timed-pregnant dams were administered a single intraperitoneal injection of 25 mg/kg EdU at embryonic day 17 (E17), and offspring were collected at postnatal day 1, paraffin embedded and sectioned. Tissues were pre-treated in 0.01 M sodium citrate buffer (pH 8) at 95°C twice for 5 minutes, then acid washed using 2 N HCl with 0.5% Triton X-100 in PBS for 1 hour at room temperature. EdU detection was performed using Click-iT kit, according to the manufacturer’s protocol (Thermo Fisher Scientific, C10337). Immunostainings were performed in combination with Edu Click-it kit staining.

## RESULTS

### The timing and location of Cre activity determine the severity of Shh pathway disruption in conditional *Smo* mutants

Smoothened (Smo) is as a critical transducer in the Shh signaling pathway, conveying signals from the Shh ligand to downstream effectors that regulate gene expression (Alcedo et al., 1996, van den Heuvel and Ingham, 1996) (Fig. 1A). To investigate the effects of Shh pathway disruption in the developing retina, we employed a conditional loss-of-function approach by using a *Smo^CKO/CKO^* allele (Long et al., 2001). This line was crossed with either Chx10-Cre or Rax-Cre drivers.

**Figure 1.**
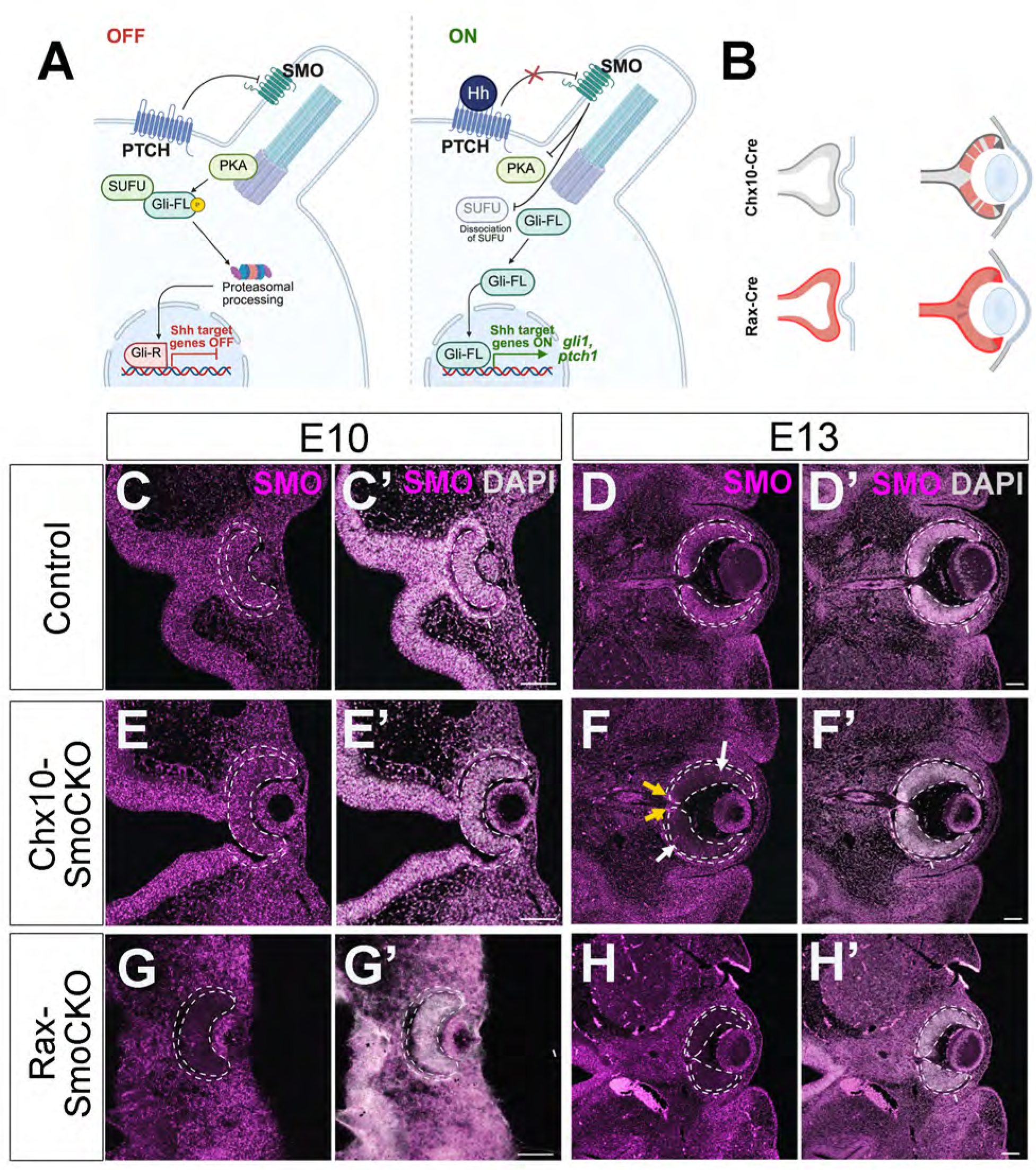
Two Cre lines drive *Smo* ablation in the developing eye. **A)** Simplified diagram of Shh signaling. **B)** The expression of Cre in Chx10-Cre and Rax-Cre lines is indicated in red, note the earlier and broader expression of Rax-Cre. **C-H)** *Smo* (magenta) expression at E10.5 and E13.5 in the different transgenic lines. Tissues have been counterstained with DAPI (gray, in C’, D’, E’, F’, G’ and H’). Note *Smo* expression in some RPCs (white arrows) and in the cells of the ONH (yellow arrows). Scale bar C-H’: 100 microns.

Chx10/Vsx2 is expressed in retinal progenitor cells (RPCs), beginning at the optic vesicle stages (Liu et al., 1994), but the onset of Chx10-Cre driver expression is later, starting at embryonic day (E) 10 and proceeding in a progressively mosaic pattern (Rowan and Cepko, 2004) (Fig. 1B). In contrast, Rax is expressed much earlier, starting at E7.5 (Furukawa et al., 1997, La Torre et al., 2015) and the Rax-Cre driver exhibits expression from at least E8. Similar to endogenous *Rax*, this transgene is expressed uniformly in the ventral thalamus/hypothalamus and optic vesicle, cup and stalk, and all subsequently arising tissues such as the RPE, ONH, ciliary body, and all retinal cell types (Bosze et al., 2020) (Fig. 1B).

Using RNAscope *in situ* hybridization (ISH) we first confirmed the removal of *Smo* mRNA in each conditional mutant (Fig. 1C-H’). As expected, in the Chx10-Cre;*Smo^CKO/CKO^* model (hereafter Chx10-SmoCKO), we did not observe changes in *Smo* expression at E10 compared to control littermates (Fig. 1C-C’, E-E’). Given the earlier and uniform activity of the Rax-Cre driver in Rax-Cre;*Smo^CKO/CKO^* (hereafter Rax-SmoCKO) optic cups, we noted the complete loss of *Smo* mRNA by E10 (Fig. 1G-G’).

At E13.5, both conditional mutants exhibited obvious loss of *Smo* mRNA expression throughout the retina. There was also a loss of *Smo* in the RPE, ONH and OS for just Rax-SmoCKO (Fig. 1H-H’), plus a concomitant reduction of the Shh direct target genes *Gli1* and *Ptch1* (Suppl. Fig. 1C-C’, F-F’). In contrast, Chx10-SmoCKO RPCs retained mosaic expression of *Smo* (Fig. 1F-F’, arrows), together with mosaic *Gli1* and *Ptch1* in RPCs, and high levels of all three transcripts in the ONH, optic stalk, RPE, and ciliary margin, outside the Chx10-Cre expression domain (Suppl. Fig. 1B-B’, E-E’, white arrows indicate clones that retained expression in the neural retina). Although both Cre drivers effectively mediate *Smo* ablation, we predicted that Rax-SmoCKO mutants will induce unique phenotypes during optic cup morphogenesis, patterning, and tissue specification, due to the earlier and broader activity of Rax-Cre. In contrast, both Rax- and Chx10-driven mutants are expected to share similar neurogenesis phenotypes. However, due to the known mosaicism of Chx10-Cre, residual wild-type cells may provide non-cell-autonomous rescue, potentially masking certain requirements for Shh signaling during retinal histogenesis.

### Rax-Cre deletion of *Smo* causes embryonic microphthalmia, coloboma, and optic nerve head defects

Because Rax-Cre is active earlier than Chx10-Cre and deletes genes uniformly in all optic vesicle derivatives (Fig. 1, and Suppl. Fig.1), we predicted that Rax-SmoCKO eyes would fully reveal roles for Shh signaling during optic cup morphogenesis. Indeed, only E13.5 Rax-SmoCKO eyes appeared smaller than those of controls. To quantify this phenotype, we measured eye diameter and length among control and Chx10-SmoCKO eyes or control and Rax-SmoCKO littermates (Fig. 2A-E). Notably, the Rax-SmoCKO eyes also uniquely had a significant reduction in vitreal space (Fig. 2F), further indicating early morphogenetic abnormalities that are not observed when *Sm*o is deleted at later stages.

**Figure 2.**
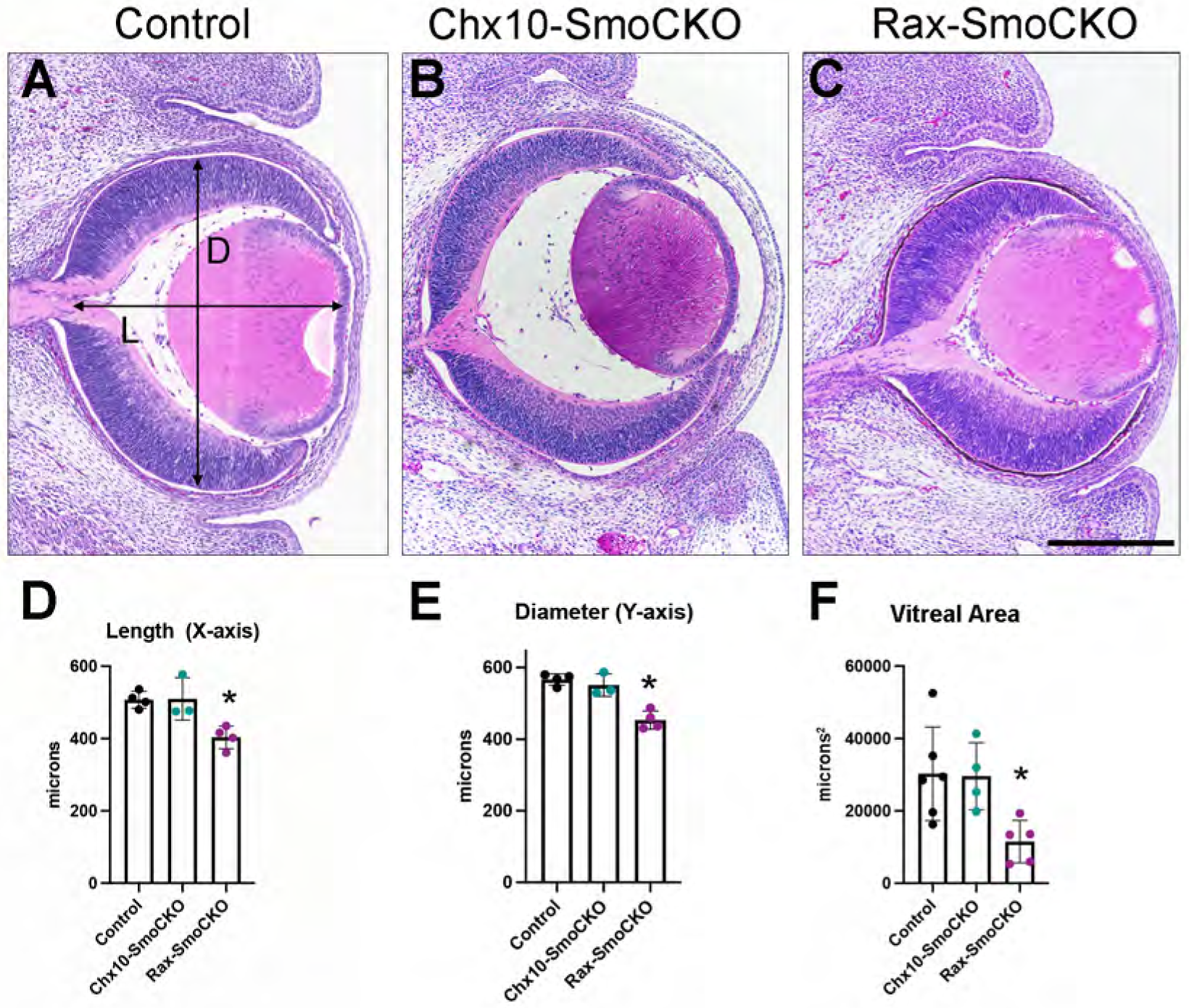
*Smo* ablation resulted in smaller eye size in Rax-SmoCKO mice, but not in Chx10-SmoCKO. **A-C)** H&E staining of paraffin sections at E13.5. Scale bar: 100 microns. **D)** Quantifications of eye length (X-axis) showed a reduction in size in Rax-SmoCKO eyes (*p*= 0.011, one-way ANOVA). **E)** Quantifications of eye diameter (Y-axis) also showed a reduction in Rax-SmoCKO eyes (*p*= 0.019, one-way ANOVA). Similarly, Rax-SmoCKO eyes exhibited a substantial decrease in vitreal area compared to the other models (*p*=0.029, one-way ANOVA).

Shh plays a critical role in ventral eye patterning (Zhao et al., 2010) and its disruption has been associated with a failure to close the optic fissure, or coloboma, in animal models (Zhang et al., 2009, Gordon et al., 2018) and human patients (Schimmenti et al., 2003). Consistent with these reports, we found that E13.5 Rax-SmoCKO eyes contained persistent optic fissures (Suppl. Fig. 2), whereas Chx10-Cre-SmoCKO eyes were indistinguishable from wild type controls (data not shown). We conclude the specificity of this conditional phenotype is likely due to the ability of Rax-Cre to remove genes from the optic vesicle, cup, and stalk whereas Chx10-Cre is confined to RPCs. Given these findings, we restricted our subsequent studies using embryonic samples to Rax-SmoCKO animals.

To further investigate how an early loss of Shh signaling impacts optic cup patterning, we examined the expression of key regional markers. In the optic vesicle, the transcription factors Pax2 and Pax6 are initially co-expressed, but soon resolve into complementary domains: Pax6 becomes restricted to the presumptive neural retina and RPE, while Pax2 localizes to the optic ONH and stalk. This spatial segregation is critical for establishing a local neural-glial boundary at the back of the retina and for distinguishing it from adjacent forebrain structures (Schwarz et al., 2000, Baumer et al., 2003, Bosze et al., 2021). Thus, we examined the expression of Pax2 and Pax6 at multiple stages of embryonic development. By E10.5, control eyes display the expected overlap of Pax2 and Pax6 in the optic vesicle, marking an early stage before the boundary between the retina and the ONH is formed (Fig. 3A-A’). In Rax-SmoCKO retinas, we also observed co-expression of these two transcription factors; however, Pax2 expression appeared reduced and more centrally restricted, suggesting a possible delay or disruption in optic stalk specification in the absence of Shh signaling (Fig. 3B-B’). By E13.5, control retinas display a well-defined boundary between the neural retina and the ONH, marked by complementary expression domains, with Pax6 confined to the retinal neuroepithelium and Pax2 restricted to the optic stalk and ONH (Fig. 3C-C’). In Rax-SmoCKO eyes, this organization is disrupted, most notably along the nasal side of the ONH (Fig.3D-D’). In many cases, Pax2+ cells were almost completely absent from the nasal ONH, and Pax6+ retinal tissue extended abnormally into the optic stalk territory, reflecting a severe disruption in the establishment or maintenance of this tissue boundary (Fig. 3G). In the temporal side, Pax6 and Pax2 were frequently co-expressed inappropriately in some cells of the ONH, and the remaining Pax2+ domain appeared narrowed and disorganized. Similarly, reductions in Pax2 expression were also observed along the ventral retina when analyzing the ventral-dorsal axis, further supporting a regionally biased disruption of optic stalk and ONH identity in the absence of Shh signaling (Fig. 3E-F).

**Figure 3.**
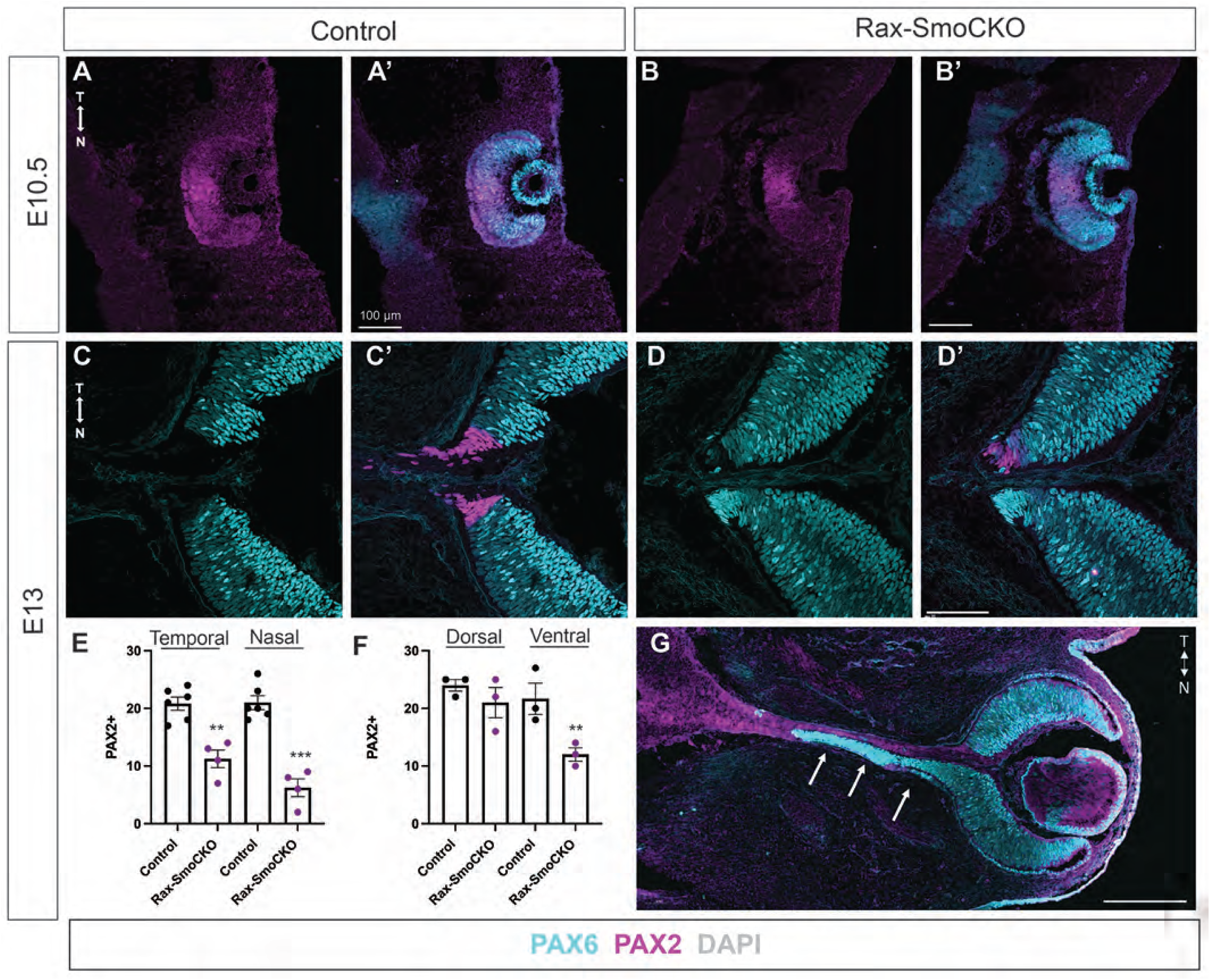
Rax-SmoCKO eyes exhibited optic nerve head defects. Pax2 (magenta) and Pax6 (teal) immunostainings at E10.5 **(A-B’)** and E13.5 **(C-D’, and G)**. **E-F)** Quantifications of the number of Pax2+ cells in the different regions of the optic cup at E13.5. Note the ectopic growth of Pax6+ retinal tissue within the nasal part of the optic cup in G (arrows). ** p<0.001, *** p<0.0005. Scale bars: 100 microns in A-B’, 50 microns in C-D’, and 150 microns in G.

### Naso-temporal patterning is affected in Rax-SmoCKO retinas

Given the unexpected ONH defects found that align to both dorso-ventral and nasal-temporal axes in Rax-SmoCKO eyes, we directly tested whether nasal-temporal retinal patterning was affected following loss of *Smo*. The establishment of this axis coincides with the mutually exclusive expression of Foxg1 in the nasal retina and Foxd1 in the temporal retina, which also act to restrict each other’s expression domains (Hatini et al., 1994, Bourguignon et al., 1998, Herrera et al., 2004, Picker et al., 2009, Takahashi et al., 2009, Carreres et al., 2011). At E10, Foxg1 and Foxd1 expression were unaffected, indicating that the initial establishment of nasal and temporal identities occurred independently of Shh signaling (Fig. 4A–B). By E13.5, however, these expression domains were disrupted in Rax-SmoCKO retinas (Fig. 4C-F): Foxg1, typically restricted to the nasal retina with a gradient that ends before the ONH, extended into the ONH domain and was ectopically expressed in temporal patches (Fig. 4D, arrows). Foxd1, normally confined to the temporal retina, also showed irregular and ectopic expression on the nasal side (Fig. 4F, arrows). These findings suggest that while Shh signaling may be dispensable for early nasal-temporal specification, it is required for maintenance and refinement of this axis at later stages.

**Figure 4.**
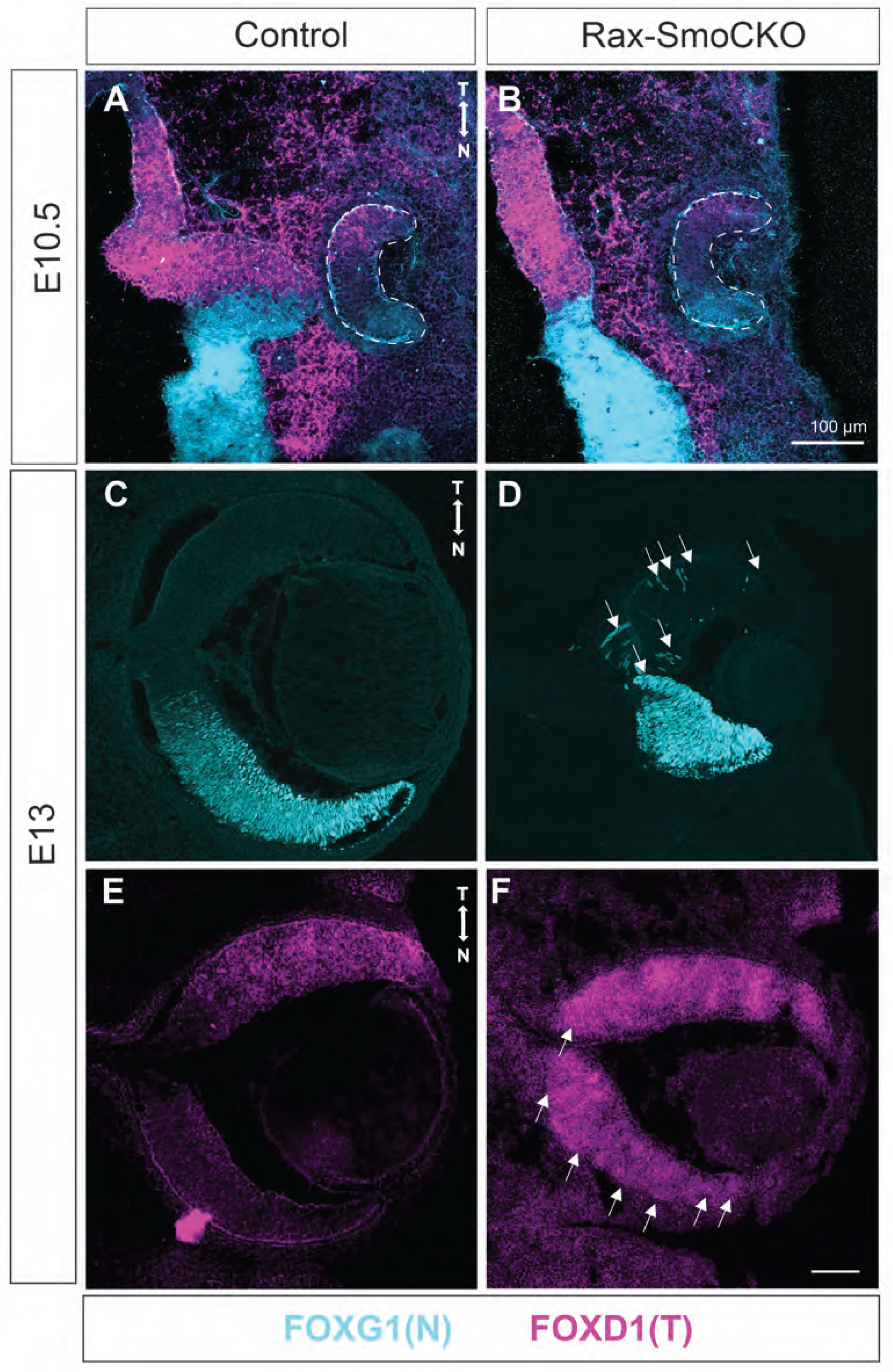
Nasal-temporal patterning defects in Rax-SmoCKO mice. FoxG1 immunostaining (teal) and FoxD1 RNA ISH (magenta) at E10.5 **(A-B’)** and E13.5 **(C-F).** Note the ectopic expression of these markers in the Rax-SmoCKO samples at E13.5 (arrows in D and F). Scale bars: 100 microns.

### At early embryonic stages, Rax-SmoCKO eyes displayed altered transcriptomic landscapes and cellular composition

Given the morphological defects observed in Rax-SmoCKO eyes, including microphthalmia, we performed bulk RNA-seq on dissected E13.5 retinas to identify underlying transcriptional changes. As expected, these experiments quantitatively validated a significant reduction in *Smo* levels in Rax-SmoCKO retinas, along with downregulation of *Gli1*, *Ptch1,* and *Ptch2* (Fig. 5A-B, Suppl. Table 1). Not all known Shh pathway targets were affected since we did not detect any significant changes in the mRNA levels of Shh co-receptor genes or genes expressed within primary cilia (Fig. 5B). Immunolabeling further confirmed the absence of significant changes in co-receptor expression, with the exception of a spatially expanded expression domain of Cdo (Suppl. Fig. 3). Interestingly, this dataset also revealed an upregulation of *Shh* itself. This increase may reflect a higher number of RGCs, the primary source of Shh in the developing retina. It has been well-characterized that as RGCs begin to differentiate, they produce and secrete Shh, which in turn, influences the proliferation and patterning of surrounding RPCs (Neumann and Nuesslein-Volhard, 2000, Zhang and Yang, 2001a, Wang et al., 2005, Ringuette et al., 2016). Therefore, elevated Shh expression could be indicative of an expanded RGC population. In support of this interpretation, several RGC transcripts such as *Pou4f1, Pou4f2, Pou4f3, Sox4, Calb*, and *Nefl* were also upregulated in the Rax-SmoCKO samples (Fig. 5A-B). A previous study (Sakagami et al., 2009) showed that the proneural bHLH factor *Atoh7*, known to regulate RGC competence (Brown et al., 2001, Brzezinski et al., 2012, Miesfeld et al., 2020), was upregulated in E15 Chx10-Cre;*Smo^CKO/CKO^*retinas. Here, we also observed an increase in *Atoh7* mRNA levels in E13.5 Rax-SmoCKO eyes (Fig. 5B). To validate this outcome, we immunostained E13.5 sections from control and Rax-SmoCKO eyes with validated Atoh7 and Brn3 (Pou4fa/b/c) antibodies (Fig. 5C-D’) and quantified marker+ cells (Fig. 5G and H, respectively). In agreement with the transcriptomic profiles, the percentages of Atoh7+ RPCs and Brn3+ RGCs were each significantly increased.

**Figure 5.**
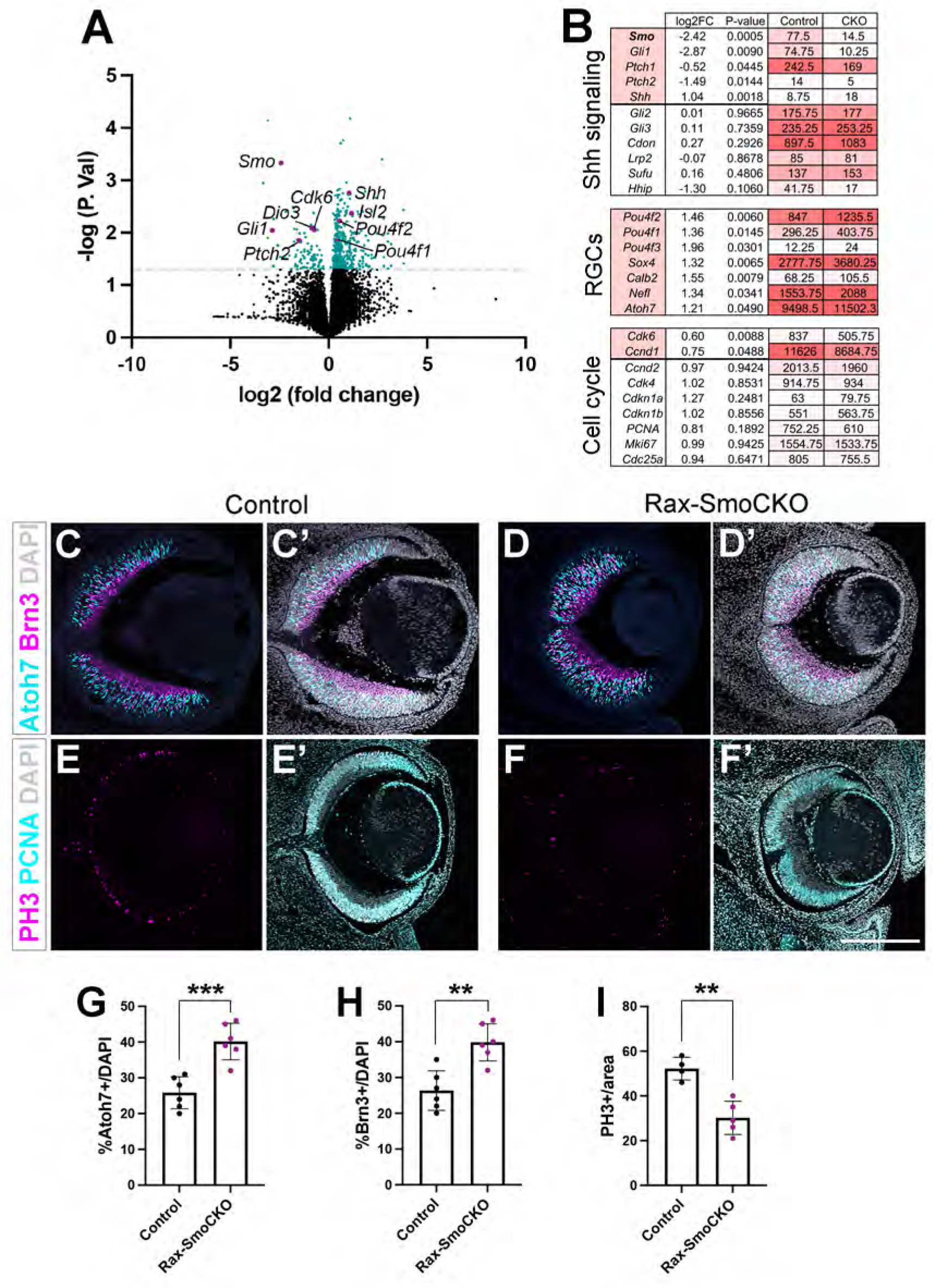
Changes in the E13.5 Rax-SmoCKO retinal transcriptome. **A)** Volcano plot highlighting some of the genes differentially expressed in Rax-SmoCKO retinas compared to control littermates at E13.5. **B)** RNAseq analyses revealed changes in Shh signaling, RGC signatures, and cell cycle genes. **C-D’)** Immunostaining with Atoh7 (teal), Brn3 (magenta) and counterstained with DAPI (gray) on control and Rax-SmoCKO E13.5 samples. E-F’) Labeling with PH3 (magenta), PCNA (teal), and DAPI (gray) in control and Rax-SmoCKO eyes. G-I) Quantifications of % of Atoh7+ (G), Brn3+ (H) cells. I) Quantification of the number of PH3+ cells normalized by area (100×10^3^ μm^2^). Scale bar: 200 microns.

In addition to changes in RGC genes, we also observed alterations in the mRNA expression of several key regulators of cell cycle progression, including *Cdk6* (Cyclin-Dependent Kinase 6) and *Ccnd1* (Cyclin D1). Both were significantly downregulated in the Rax-SmoCKO samples (Fig, 5B, Suppl. Table 1). We further validated the reduction in *Cdk6* by ISH (Suppl. Fig. 4). Cdk6 and Cyclin D1 form a complex to regulate the G1 phase of the cell cycle as well as G1-to-S transition (Malumbres, 2014, Bury et al., 2021). Thus, the changes observed suggest a loss of RPC proliferative capacity at this stage. To test this possibility, we labeled E13.5 retinas with anti-phospho-Histone H3 (PH3), a well-established marker of actively dividing cells. This further confirmed a significant reduction of cells in mitosis in Rax-SmoCKO retinas, compared to controls (Fig. 5E–F’, I).

Notably, other molecules were also altered in Rax-SmoCKO eyes, including type 3 iodothyronine deiodinase (*Dio3),* an enzyme responsible for degrading thyroid hormone (TH). TH signaling is known to play a critical role in regulating cone photoreceptor maturation and subtype specification (Ng et al., 2010, Eldred et al., 2018, McNerney and Johnston, 2021). For instance, loss of TH signaling leads to a reduction or absence of L/M cones, whereas exposing human retinal organoids to high levels of T3 results in a marked increase in L/M cone density (Eldred et al., 2018). We noted a loss of *Dio3* mRNA in the absence of Smo activity, which was validated by ISH (Suppl. Fig. 4B-B’, D-D’). Given the change in *Dio3* expression observed, we investigated whether cone subtype specification was altered in Rax-SmoCKO retinas. For these experiments, we stained P21 flat-mounted retinas with S-Opsin and M-Opsin antibodies. Notably, Rax-SmoCKO retinas were smaller and contained an overall higher concentration of cone photoreceptors, but the normal enrichment of true S-cones in the ventral side of the retina (Nadal-Nicolas et al., 2020) was unaffected (Suppl. Fig. 5A-F). There were no significant changes in the proportion of different cone subtypes in the dorsal retina (S-Opsin+ M-Opsin-cells, *p*=0.27 vs control; S-Opsin-M-opsin+ *p*= 0.18 vs control; and S-Opsin+ M-Opsin+, *p*=0.38 vs control, Suppl. Fig 5G). In contrast, the ventral retina showed significant alterations, including an increase in true S-cones (S-Opsin+ M-Opsin-, *p*=0.014) concomitant with a reduction in mixed S-Opsin+ M-Opsin+ cones (p=0.007), but no changes in S-opsin-M-Opsin+ cones (*p*=0.34, Suppl. Fig 5G).

### Postnatal ocular defects are more severe following early Smo ablation with Rax-Cre compared to Chx10-Cre

Since microphthalmia was already evident at E13.5, we assessed eye morphology at postnatal day 21 (P21) to evaluate the sustained effects of Smo ablation using either Rax-Cre or Chx10-Cre drivers. Grossly, Chx10-SmoCKO mutant eyes are indistinguishable from controls but Rax-SmoCKO mutants have microphthalmic eyes (Fig. 6A-C). Histologic sections of Chx10-SmoKO eyes reveal that both the outer nuclear layer (ONL) and inner nuclear layer (INL) were significantly thinner (*p*= 0.0007 and *p*<0.0001, respectively, Fig. 6D-E, G-H, J). At the same time, no significant changes were observed in the overall thickness of the ganglion cell layer (GCL) (Fig. 6J, *p*= 0.63), although there was a significant increase in the number of cells within this layer (Fig. 6K, *p*= 0.038). The rest of the eye, namely the lens, ciliary body, and anterior chamber were not different from the controls (Fig. 6B, E). In sections of microphthalmic Rax-SmoCKO eyes, we observed less vitreal space, and the ciliary bodies seemed to be missing (Fig. 6C, F, and Suppl. Fig. 6). There was also extensive thinning of the ONL and INL (Fig. 6F, I, K; both p <0.0001), accompanied by a dramatic expansion of the GCL, with significantly more cells (Fig. 6D-I, K, Control vs Rax-SmoCKO p<0.0001, Chx10-SmoCKO vs Rax-SmoCKO p=0.0005). Rax-SmoCKO eyes also had abnormal anterior chambers (Fig. 6F) and aberrant lenses, with possible duplication and reorientation of the anterior epithelial layer (Suppl. Fig. 6), even though there is no Cre expression associated with these tissues. Consequently, these anterior segment abnormalities likely result from non-cell autonomous effects of early retinal malformation.

**Figure 6.**
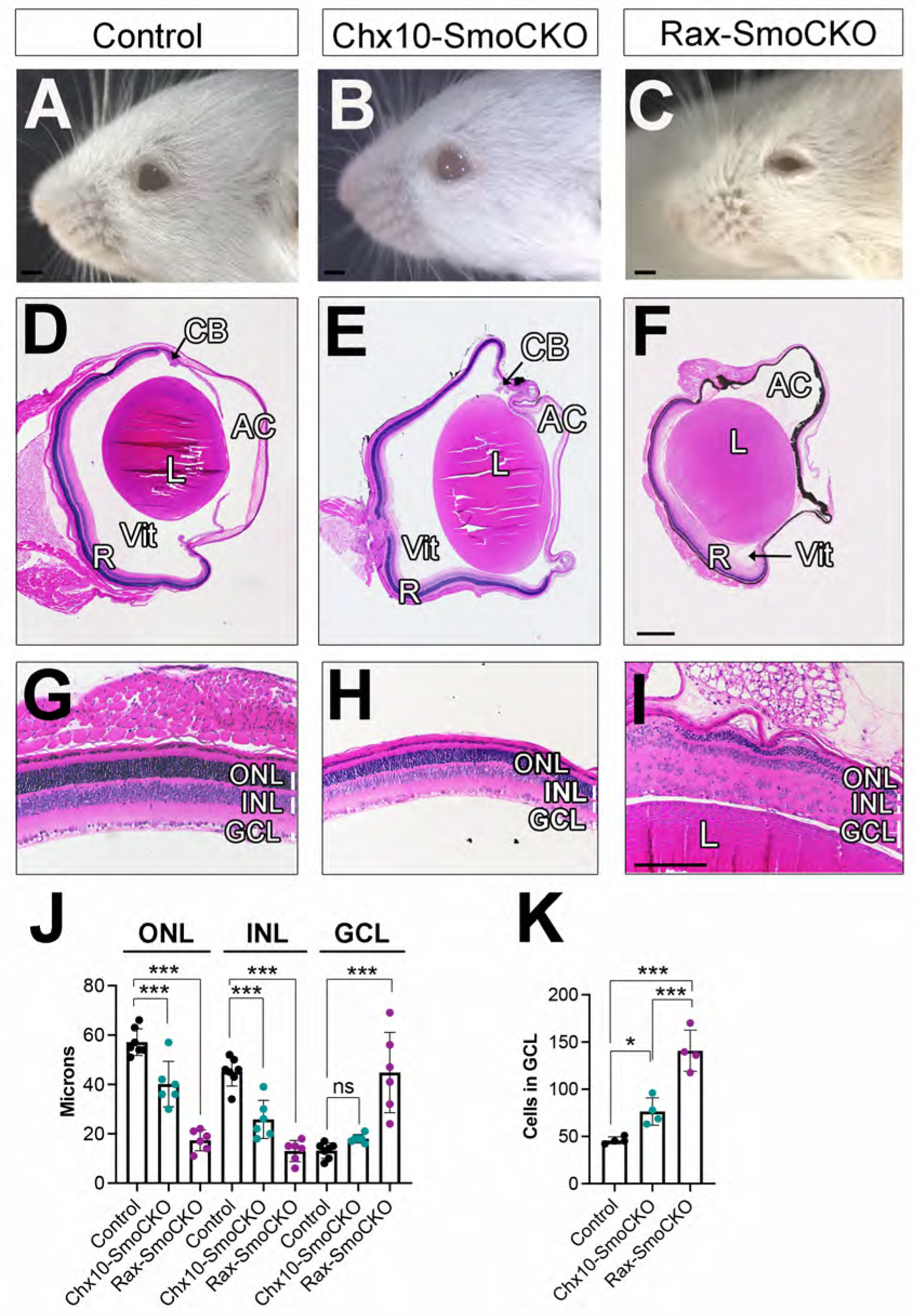
Adult Chx10-SmoCKO and Rax-SmoCKO eye phenotypes. **A-C)** At P21, only Rax-SmoCKO mice show gross morphological defects with “squinting” eye appearance, consistent with microphthalmia and/or anterior segment dysgenesis. **D-I)** H&E staining on paraffin sections showed retinal thinning in both the Chx10-SmoCKO and Rax-SmoCKO models. **J)** Quantifications of layer thickness in microns. **K)** Quantification of the number of nuclei in the GCL (in 500 microns). One-way ANOVA multiple comparison tests were used to obtain p-values: * p<0.005, *** p<0.0005, **** p< 0.0001. Scale bars: 5mm in A-C. 250 microns in D-F, 100 microns in G-I.

### Sonic hedgehog signaling influences cell population ratios in the retina

All retinal neurons and the Müller glia are born from a single population of multipotent RPCs following a conserved neurogenic program (Cepko et al., 1996, Livesey and Cepko, 2001, Bassett and Wallace, 2012, Clark et al., 2019, Lu et al., 2020). Cell fate specification is accomplished through a network of both intrinsic and extrinsic factors (Kim et al., 2005, Emerson et al., 2013, La Torre et al., 2013, Mattar et al., 2015, Zhang et al., 2023a, Javed et al., 2023), and Shh signaling is known to be required by RPCs in a cell-autonomous manner to maintain the progenitor pool (Jensen and Wallace, 1997, Levine et al., 1997, Black et al., 2003, Moshiri et al., 2005, Wang et al., 2005), with Smo loss leading to a substantial increase in RGC numbers (Wang et al., 2005, Sakagami et al., 2009). Consistent with this, our data using the Rax-Cre driver demonstrated increased numbers of RGCs at E13.5, aligning with prior observations in the Chx10-SmoCKO retina, which showed elevated RGC numbers at E15 and P0 (Sakagami et al., 2009). A mild increase in Otx2+ Crx+ photoreceptors was also reported at E15.5 in the Chx10 model. However, because retinal neurogenesis is ongoing until postnatal P7, these findings may represent a partial view of the consequences of *Smo* ablation throughout retinal development. To gain a more comprehensive understanding, we took advantage of the uniformly expressed Rax-Cre driver to investigate other effects of *Smo* deletion at P21, after retinogenesis is complete.

We quantified how *Smo* ablation affects the relative abundance of multiple retinal cell types in mature retinas. Rax-SmoCKO mice not only showed an increased ratio of RBPMS+ Tuj1+ RGCs (83 vs 23 RGCs/area (100×10^3^ ^m2) in control littermates, *p*= 0.0013, Fig 7A-C), but also exhibited a uniform increase in all early-born populations. Consistent with our flat-mounted data (Suppl. Fig. 5), we found that Rax-SmoCKO retinas contained higher concentrations of cone photoreceptors (Arr3+ Otx2+, 63.33 vs 23.67 cones/area in controls, *p*= 0.009, Fig. 7D-F), along with an increase in horizontal cells (Calbindin+, 10.25 vs 3.6/area in control retinas, *p*= 0.0072. Fig. 8A-C), and cholinergic amacrine cells (starburst amacrine cells, SACs, 31.15 vs 11.3 SACs/area in control retinas *p*= 0.022). SACs can be identified by Sox2 labeling (Whitney et al., 2014), but Sox2 is also expressed by Müller glia and retinal astrocytes, potentially clouding the effect on SACs specifically. To get around this issue, we co-labeled retinal sections with Sox9, a known marker of Müller glia and astrocytes, and quantified each single- and double-labeled population, with Sox2+ Sox9-representing the SACs (Fig. 8D-F).

**Figure 7.**
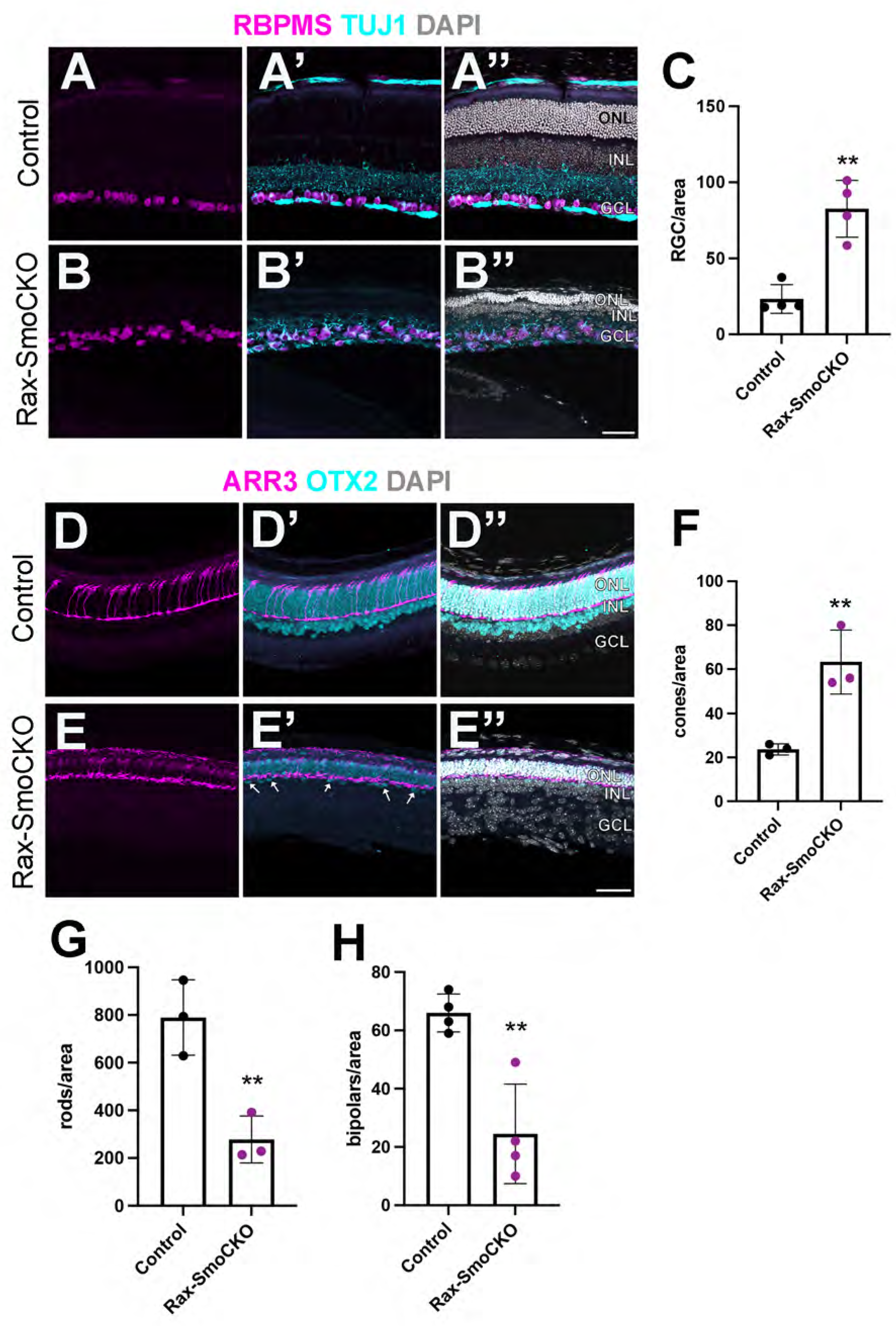
P21 Rax-SmoCKO retinas have more RGCs and cones, at the expense of rods and bipolar cells. **A-B’’)** RBPMS (magenta) and Tuj1 (teal) in control retinas (A-A’’) and Rax-SmoCKO (B-B’’). **C)** Quantifications of the number of RGCs, as represented by RBPMS+ Tuj1+ cells in control and Rax-SmoCKO retinas. Cell numbers are normalized by area (100×10^3^ μm^2^). **D-E’’)** Cone Arrestin (Arr3, magenta) and Otx2 (teal) stainings in the different lines. The arrows in E’ indicate the small number of Otx2+ bipolar cells in Rax-SmoCKO. **F)** Quantification of cone photoreceptors, as represented by the number of Cone Arrestin+ Otx2+ cells normalized by area. All samples were counterstained with DAPI (gray). **G)** Quantification of the number of rods, as represented by the number of Otx2+ Cone Arrestin-cells in the ONL. **H)** Quantification of bipolar cells, measured by the number of Otx2+ cells in the INL. ONL: outer nuclear layer, INL: inner nuclear layer, GCL: ganglion cell layer. * p<0.005, ** p<0.001. Scale bar: 100 microns.

**Figure 8.**
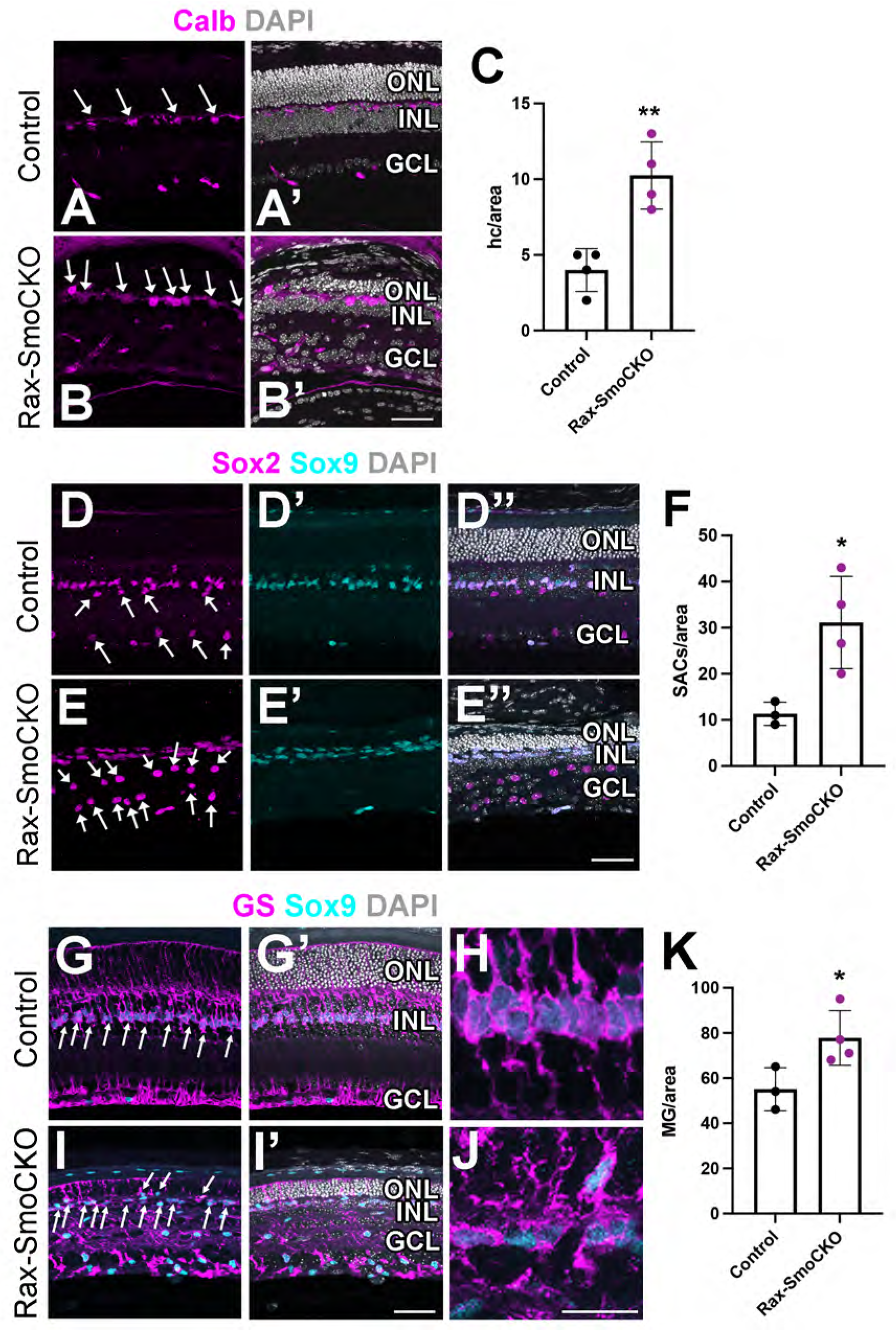
Rax-SmoCKO adult retinas also exhibit higher ratios of horizontal cells, starburst amacrine cells, and Müller glia. **A-B’)** Calbindin (Calb, magenta) stainings in Rax-SmoCKO and control retinas. Samples were counterstained with DAPI (gray, A’ and B’). **C)** Quantification of the number of Calb+ horizontal cells in control and Rax-SmoCKO retinas. Cell numbers are normalized by area (100×10^3^ μm^2^). **D-E’’)** Sox2 (magenta), Sox9 (teal), and DAPI (gray) labeling in the different lines. The arrows in D and E indicate Sox2+ Sox9-starburst amacrine cells. Also, note the expansion in Sox9+ cells in the INL. **F)** Quantification of starburst amacrine cells, as represented by the number of Sox2+ Sox9-cells normalized by area. **G-J)** Labeling with Glutamine Synthetase (GS, magenta), Sox9 (teal) and DAPI (gray). Note the increase in Müller glia cells (arrows) in I-I’. **H and J**) show close-ups of Müller glia cells in the different mouse lines. **K)** Quantification of Müller glia cells, as represented by the number of GS+ Sox9+ cells normalized by area. ONL: outer nuclear layer, INL: inner nuclear layer, GCL: ganglion cell layer. * p<0.005, ** p<0.001. Scale bar: 100 microns in A-I’ and 20 microns in H and J.

Notably, cone photoreceptors displayed altered morphologies with shortened axons and outer segments (Fig. 7E), indicating potential disruptions in their structural development.

In contrast to the expansion of early-born populations, late-born retinal cell types were significantly reduced. In this direction, Rax-SmoCKO retinas also showed a significant reduction in rod photoreceptors (Otx2+ Arr3-cells in the ONL, 278 rods vs 789.7 rods/area in controls, *p*= 0.009, Fig. 7D-E’’, G) and bipolar cells (Otx2+ in the INL, 24.5 vs 66 bipolar cells/area in controls, *p*= 0.0039, Fig. 7D-E’’, H). Because the shift toward early-born neurons occurred at the expense of later-born types, these phenotypes suggested altered temporal fate specification. Interestingly, Müller glia did not follow this pattern. Instead, we observed an increase in the number of Sox9+ GS+ Müller glia cells (77.75 vs 55 cells/area in controls, p-value: 0.044, Fig. 8G-K). Some of these were ectopically positioned in the ONL and displayed signs of abnormal polarity (Fig. 8J).

It has been proposed that RGCs help orchestrate the spatial and temporal patterning of astrocytic invasion from the ONH, which spreads radially across the retina during development. In turn, astrocytes support vascular growth by providing structural integrity and releasing angiogenic signals (Edwards et al., 2012, O’Sullivan et al., 2017). This interplay between RGCs, astrocytes, and retinal vasculature is crucial for the proper formation of the retinal microenvironment. Given the dramatic increase in RGCs in Rax-SmoCKO, we also assessed the indirect impacts on astrocytes and vascular development. As shown in Suppl. Fig. 7, *Smo* loss led to an important increase in astrocyte cell numbers. In control conditions, each retinal astrocyte contacts at least one blood vessel, and extends end-feet that surround the vessels, forming a close structural relationship with the vascular wall. Instead, in Rax-SmoCKO retinas, astrocytes displayed flat almost “carpet-like” morphologies that in central regions covered the whole surface of the retina. Consistently, we also observed aberrant blood vessel morphologies (Suppl. Fig. 7).

### Loss of Smo activity directly impacts temporal retinal cell competence

Overall, our data suggest that, in the absence of *Smo*, RPCs exit the cell cycle prematurely, potentially leading to an overproduction of early-born cell types at the expense of later-born populations. An alternative, but not mutually exclusive, possibility is that Shh signaling could also play a direct role regulating RPC competence (*i.e.,* the ability of progenitors to generate specific retinal cell types at different times of development) rather than solely influencing their timing of cell cycle exit.

In order to assess whether RPC competence was affected in Rax-SmoCKO retinas, we labeled progenitors with 5-ethynyl-2’-deoxyuridine (EdU) at E17, a stage where the progenitors in the central retina have normally transitioned to a late competence stage and have lost the ability to generate early cell types such as RGCs and cone photoreceptors (Young, 1985, Reese and Colello, 1992, Brzezinski et al., 2012, Clark et al., 2019). Retinas were then harvested at P1, and we assessed the fate identities of the EdU-labeled cells in the central retina. As expected, EdU+ RBPMS+ RGCs were absent in control retinas (Fig 9). Strikingly, however, numerous EdU+ RBPMS+ cells were observed in Rax-SmoCKO eyes (*p*=0.007), along with a significant increase in EdU+ S-opsin+ cones (*p*=0.0008, Fig. 9). These findings indicate that loss of Smo extends the window of early progenitor competence, allowing for the continued production of early-born retinal neurons beyond their normal developmental timeframe.

**Figure 9:**
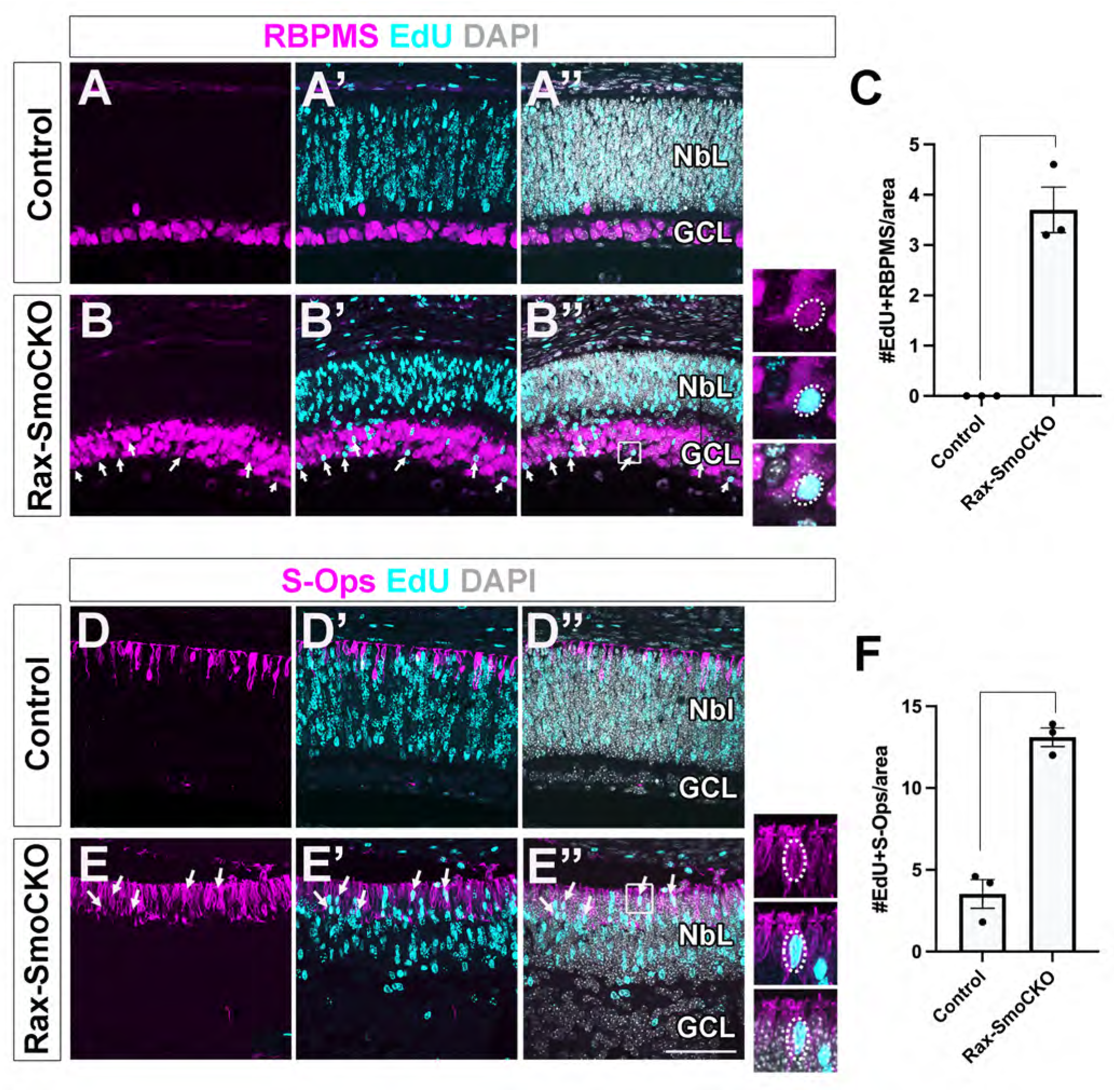
RGCs and cone photoreceptors were generated at later times in Rax-SmoCKO retinas. Control and Rax-SmoCKO mice were injected with EdU at E17 to label RPCs and samples were collected at P1. **A-B’’)** Co-staining with RBPMS (magenta), EdU (teal), and DAPI (gray) revealed EdU+ RPMS+ cells in the central retinas of Rax-SmoCKO mice (arrows) but not in controls. **C)** Quantifications of the number of EdU+ RBPMS+ cells normalized by area. **D-E’’)** Co-staining with S-Opsin (magenta), EdU (teal), and DAPI (gray). Arrows show EdU+ S-Opsin+ cells. **F)** Quantification of EdU+ S-Opsin+ cells, normalized by area. NbL: Neuroblastic layer, GCL: Ganglion Cell layer. Scale bar: 100 microns.

## DISCUSSION

This study uncovers previously unrecognized requirements for Shh signaling in the mammalian eye, revealed through the targeted disruption of *Smo* using a Cre driver that uniformly deletes genes among all optic vesicle derivatives and across all stages of retinal development. While Shh is already known as key player during early optic vesicle morphogenesis (Chiang et al., 1996, Zhang and Yang, 2001b) and progenitor cell proliferation (Jensen and Wallace, 1997, Levine et al., 1997, Moshiri et al., 2005, Wang et al., 2005, Wall et al., 2009, Bibliowicz and Gross, 2011), our findings highlight additional key functions during patterning and cell fate specification. Our experiments also revealed distinct alterations depending on the timing and location of *Smo* ablation. Thus, while Chx10-Cre-driven deletion caused modest changes limited to the retina, Rax-Cre-driven deletion led to broader defects that affect the retina, ONH, ciliary body, lens, and anterior chamber. These differences reflect the wider expression domain of Rax-Cre, but also uncover non-cell autonomous effects, as, for instance, the lens and anterior chamber are not within the Rax-Cre domain. These findings are also consistent with similar outcomes in Lhx2 conditional mutants (Yun et al., 2009, Thein et al., 2016), where Lhx2 deletions using several Cre drivers with different tissue specificities led to loss of retinal structures along with indirect defects in the lens and anterior segment due to disrupted BMP and FGF signaling, supporting the idea that early retinal perturbations can influence the surrounding tissues.

The more pronounced defects in the Rax-SmoCKO mutants are also likely the result of its earlier onset of expression compared to Chx10-Cre (Furukawa et al., 1997). Similarly, the mosaic nature of the Chx10-Cre driver (Rowan and Cepko, 2004) could partially obscure the full extent of retinal phenotypes and contribute to the milder effects observed. These findings underscore the critical importance of both the spatial and temporal regulation of Shh signaling to coordinate multiple aspects of eye development.

Among the more severe consequences of early and widespread *Smo* ablation in the Rax-SmoCKO mutants were prominent defects in early eye patterning. One of the most evident abnormalities was the presence of coloboma, consistent with a failure in optic fissure closure. This phenotype is well-aligned with the established role of Shh signaling in ventral eye development, as disruption of this pathway commonly results in coloboma across vertebrate models and in human patients (Schimmenti et al., 2003, Morcillo et al., 2006, Lee et al., 2008, Gordon et al., 2018).

Notably, additional patterning defects were observed in the ONH, including an apparent reduction in Pax2 expression and a failure to properly delineate the ONH domain. This disruption is especially evident on the nasal and ventral portions of the eye. In some samples, this defects in ONH result in retinal tissue inappropriately extending into the nasal optic stalk territory. Pax2 expression is already diminished as early as E10, suggesting that Shh signaling is required early in development to establish proper ONH identity. These findings support previous work showing that Pax2 is a downstream target of Shh (Ekker et al., 1995, Macdonald et al., 1995) and that Shh secreted by RGCs is essential for maintaining Pax2 expression and supporting glial development in the optic nerve (Dakubo et al., 2003). Our data build on this model and suggest that Shh signaling also plays an earlier role in ONH patterning, potentially by coordinating regional identity between the optic stalk and adjacent retinal tissue. This highlights a broader requirement for Shh in maintaining spatial boundaries and organizing the architecture of the developing eye.

Similarly, our findings indicate that Shh signaling is critical for maintaining nasal-temporal patterning in the retina, but not for its initial establishment. At E10, no disruptions in nasal-temporal patterning were observed, but by E13.5, we observed ectopic expression of both FoxG1 and FoxD1, highlighting a role for Shh in sustaining these identities. This contrasts with studies in zebrafish, where Shh signaling is important for the establishment of nasal-temporal patterning (Hernandez-Bejarano et al., 2015). Notably, in fish, Shh and FoxD1 identity have been shown to play a crucial role in determining the location of the acuity zone (Hernandez-Bejarano et al., 2022), an area analogous to the human fovea (Yoshimatsu et al., 2020, Lahne, 2023). And recently, missense Shh mutations have been identified in patients with abnormally positioned foveae (Azuma et al., 2025), further emphasizing the relevance of Shh signaling in shaping the spatial patterning and organization of the mammalian retina.

In line with previous findings using the Chx10-Cre model (Sakagami et al., 2009), we observed a significant increase in the number of RGCs following *Smo* ablation. However, we also identified more global changes in cell type proportions, including increased numbers of cone photoreceptors, horizontal cells, and early-born amacrine cells, accompanied by a reduction in rods and bipolar cells. The Sakagami study reported only a modest increase in cones and no significant changes in horizontal or amacrine populations. These differences may result from the developmental time points examined or the markers used to define specific cell types. For instance, the different amacrine phenotypes could stem from variations in the responses of different amacrine subtypes, as our data suggest an expansion of early-born GABAergic amacrine cells that may not have been captured in broader analyses of all amacrine populations. Our analyses of *Smo*-depleted retinas at embryonic stages revealed reduced mitotic activity, consistent with previous work in different species showing that Shh promotes retinal progenitor proliferation and delays their exit from the cell cycle (Jensen and Wallace, 1997, Wang et al., 2005, Wallace, 2008, Sakagami et al., 2009). Taken together, our findings support the hypothesis that *Smo* ablation caused RPCs to exit the cell cycle precociously, leading them to adopt fates available at that time, resulting in an overproduction of early-born cell types. However, in addition to promoting premature cell cycle exit, Shh loss-of-function also altered the competence state of RPCs. In this direction, we observed an unexpected increase in RGC and cone photoreceptor production at later stages of development, indicating that, in the absence of Shh signaling, RPCs retain the ability to generate early-born cell types beyond their typical developmental window. This shift in temporal competence reveals that Shh is not only important for the maintenance of the progenitor pool but also help orchestrate the progression of intrinsic competence states over time. These findings are consistent with a growing body of literature implicating the transcription factor Lhx2 as a key regulator of progenitor competence and neurogenic output. For example, several studies have shown that Lhx2 ablation disrupts the orderly transition of competence states, in part by altering the chromatin landscape of RPCs (Gordon et al., 2013, Zibetti et al., 2019, de Melo et al., 2016a, de Melo et al., 2016b). Importantly, Lhx2 has also been identified as an intrinsic modulator of Shh signaling during early retinal development (Li et al., 2022) suggesting a relationship between Lhx2 and Shh in coordinating progenitor maintenance and fate specification. Our findings support a model in which Shh signaling acts downstream of Lhx2 to regulate both the timing of cell cycle exit and the dynamic progression of RPC competence, thereby ensuring proper temporal patterning of retinal cell types.

In addition to an overall increase in cone photoreceptors, we also observed alterations in cone subtypes, with a significant upregulation in the number of S-cones in the ventral retina of Rax-SmoCKO mice. Thyroid hormone (TH) signaling is an essential regulator of cone subtype identity, with Dio3 contributing to S-cone specification (McNerney, 2025, Ng et al., 2010, Eldred et al., 2018). Consequently, the downregulation of *Dio3* observed in our RNA-seq data would be expected to increase T3 levels and reduce the number of S-cones. Instead, we found a ventrally restricted increase in S-cones, suggesting that TH signaling alone does not fully explain this outcome. One possible explanation is that premature cell cycle exit, caused by loss of Shh signaling, biases progenitors towards early-born fates, consistent with the time of differentiation, even within the cone lineage. Alternatively, the selective enrichment of S-cones in the ventral retina may reflect a disruption of regional identity due to impaired Shh signaling, which is known to play a critical role in establishing ventral retinal characteristics. In this context, the increase in ventral S-cones could represent a direct consequence of altered patterning cues, or a compensatory response to the failure to properly specify ventral retinal identity during early development.

Finally, we also observed changes in retinal glial populations, including astrocytes. These findings further confirm a body of evidence that RGCs are required for the migration of retinal astrocytes, which in turn guide the migration and patterning of the adult retinal vasculature (Fruttiger et al., 1996, Burne and Raff, 1997, Wallace and Raff, 1999, Edwards et al., 2012, Paisley and Kay, 2021, O’Sullivan et al., 2017). Thus, the increased number and altered distribution of RGCs we observe following Shh pathway disruption subsequently and indirectly influence astrocyte patterning and vascular development.

Taken together, our findings reveal that Shh signaling influences multiple aspects of retinal development, including progenitor maintenance, neurogenesis, cell fate specification, and spatial patterning. The wide range of phenotypes observed following *Smo* ablation, from altered cell cycle progression and shifts in neuronal subtype identity to changes in tissue patterning, highlights the sensitivity of the developing eye to disruptions in this pathway. These results are especially relevant in the context of human disorders such as Curry-Jones syndrome, which is caused by mosaic activating mutations in *Smo* and is associated with ocular abnormalities, including coloboma and microphthalmia (Twigg et al., 2016). Our data offer a developmental framework to better understand how misregulation of Shh signaling may contribute to the eye defects seen in such conditions. More broadly, this work emphasizes the essential role of Shh in coordinating the development of multiple retinal lineages and ensuring the proper organization of eye structure.

## Acknowledgements

We thank Sarah Kiser for technical assistance and all members of the Brown, Simó, and La Torre laboratories for their helpful insights. We also thank Drs. Tom Glaser and Nick Marsh-Armstrong for valuable comments and generosity with reagents. This study was supported by R01 EY013612 to NLB, EY031724 to ALT and NLB, and R01 EY013760 to EML. We benefited from the use of the National Eye Institute Core Facility for histology sample processing [supported by P30 EY012576]. The sequencing library preparations and the sequencing were carried out at the UC Davis Genome Center DNA Technologies and Expression Analysis Core, supported by NIH Shared Instrumentation Grant 1S10OD010786-01 and analyzed by the UC Davis Bioinformatics Core.

## Supplementary Figures

**Suppl. Figure 1. Downregulation of *Gli1* and *Ptch1* upon *Smo* ablation.**

**A-C’)** RNAscope in situ hybridization of *Gli1* (magenta). **D-F’)** RNAscope in situ hybridization of *Ptch1* (magenta). In all cases, the tissue was counterstained with DAPI (gray, A’. B’. C’, E’, E’, F’). Note the mosaicism of the Chx10-SmoCKO model reflected in patches of *Gli1*+ and *Ptch1*+ in RPCs (white arrows) and high levels of both *Gli1* and *Ptch1* in the ONH and optic stalk (yellow arrows). Scale bar: 100 microns

**Suppl. Figure 2. Only Rax-SmoCKO mice exhibit coloboma**

**A-B)** Photomicrographs of control and Rax-SmoCKO mice at E13.5. Note the coloboma in B (arrow). **C-D)** Sagital sections of E13.5 samples stained with Laminin (magenta) and DAPI (gray). Note the gap in the ventral part of the optic cup in in D (arrow). Scale bars: 1.5mm in A-B, 100 microns in C-D. For these experiments, we examined n = 12 Rax-SmoCKO mutants, n =7 Chx-SmoCKO mutants and n=17 WT eyes.

**Suppl. Figure 3. *Smo* ablation did not effect primary cilia position, nor Shh co-receptor expression.**

Immunostaining for Arl13b (teal), a protein enriched in primary cilia, and the RGC marker Brn3 (magenta, **A-B**). The expression of Shh co-receptors were directly compared by labeling with Lrp2 (teal) and Cdo (magenta, **C-D**), Lrp2 (teal) and Gas1 (magenta, **E-F**), and Lrp2 (teal) and Boc (magenta, **G-H**) in E13.5 eyes. Scale bar: 200 microns.

**Suppl. Figure 4. *Smo* ablation reduced Dio3 and Cdk6 mRNA expression in E13.5 retinas.** *In situ* hybridization (RNAscope) for *Cdk6* (teal, **A-A’ and C-C’**) and *Dio3* (magenta, **B-B’ and D-D’**) are shown. All the samples were counterstained with DAPI (gray). Scale bar: 100 microns.

**Suppl. Figure 5: At P21, Rax-SmoCKO retinas exhibited higher ratios of cone photoreceptor cells and changes in the proportions of cone subtypes in the ventral retina. A-B’’)** Flat-mounted retinas stained with Blue-Opsin (S-Ops, teal) and Red/green-Opsin (M-Ops, red). **C-F)** Close-up images of the outer segments in the different regions. **G)** Quantifications of the ratios of cone subtypes, classified as shown in (Nadal-Nicolas et al., 2020). Note the increase in true S-cones in the ventral region of the Rax-SmoCKO retina. Scale bars: 100 microns for A-B’’, 50 microns for C-F

**Suppl. Figure 6. Rax-SmoCKO mice have defective lenses and lack ciliary bodies.**

**A-C)** H&E staining of the different mouse lines showing the presumptive absence of ciliary body in Rax-SmoCKO eyes but not in Chx10-SmoCKO. **D-E)** DAPI staining of control and Rax-SmoCKO lens. R: retina, CB; ciliary body, L: lens. Scale bars: 100 microns in A-C, 250 microns in D-E.

**Suppl. Figure 7. Rax-SmoCKO mice exhibited defects in retinal astrocytes and retinal vasculature.** P21 flat-mounted retinas were stained with GFAP (red) and Isolectin-IB4 (teal). ONH: optic nerve head. Scale bars: 100 microns.

**Suppl. Table 1: RNA-Sequencing expression data from control and Rax-SmoCKO E13.5 eyes.**

Normalized RNA-seq counts for genes detected in four control (CONTROL1-4) and four Rax-SmoCKO (SMO1-4) samples.

## Notes

### Competing Interest Statement

The authors have declared no competing interest.

